# Mixture Margin Random-effects Copula Models for Inferring Temporally Conserved Microbial Co-variation Networks from Longitudinal Data

**DOI:** 10.1101/2022.04.25.489333

**Authors:** Rebecca A. Deek, Hongzhe Li

## Abstract

Longitudinal microbiome studies, in which data on a single subject is collected repeatedly over time, are becoming increasingly common in biomedical research. Such studies provide an opportunity to study the inherently dynamic nature of a microbiome in a way that cannot be done using cross-sectional studies. In this paper, we develop random-effects copula models with mixed zero-beta margins to identify biologically meaningful temporally conserved co-variation between two bacterial taxa, while accounting for the excessive zeros seen in 16S rRNA and metagenomic sequencing data. The model assumes a random-effects model for the dependence parameter in the copulas, which captures the conserved microbial co-variation while allowing for a time-specific dependence parameters. We develop a Monte Carlo EM algorithm for efficient estimation of model parameters and a corresponding Monte Carlo likelihood ratio test for the mean dependence parameter. Simulation studies show that our test controls the Type I error rate and provides an unbiased estimate of the mean dependence parameter. Additionally, we apply our method to a longitudinal pediatric cohort and identify changes in both local and global patterns of microbial co-variation networks in infants treated with antibiotics. Our analysis shows that the no antibiotics network is less dependent on individual taxon, thus making it more stable than the antibiotics network and more robust to both targeted and random attacks.

**Author summary:** Identification of co-variation between two microbes in microbial communities provides important insights into the community structure and stability. The commonly used measures of co-variation do not handle excessive zeros observed in the data and cannot be applied to longitudinal microbiome data directly. In this paper, we develop random-effects copula models with mixed zero-beta margins to identify biologically meaningful temporally conserved co-variation between two bacterial taxa, while accounting for the excessive zeros seen in 16S rRNA and metagenomic sequencing data. The model captures the conserved microbial co-variations while allowing for a time-specific dependence parameters. We develop an efficient Monte Carlo-based algorithm for parameter estimation and statistical inference. We analyze the data from a pediatric longitudinal cohort and identify changes in both local and global patterns of microbial co-variation networks in infants treated with antibiotics.

## Introduction

A microbiome, and its set of interactions, form a complex and dynamic ecosystem [1, 2]. Such systems are emergent; they are characterized by properties that arise from the interactions between parts, but not by any individual component [3]. It is useful to measure these emergent properties over time since it is likely that their short- and long-term effects differ. The rise in longitudinal microbial sequencing studies provides such an opportunity, as they allow for the examination of the natural variability of a microbiome. This is in contrast to cross-sectional studies, which have been successful in microbial diversity and differential abundance analyses [4–7]. Though, they are limited in that they only provide a snapshot of microbial dynamics at a single point in time. Longitudinal studies allow for full reconstruction of dynamics as a function of time.

Longitudinal microbiome data allows us to identify temporally conserved microbial co-variation networks while allowing for microbial compositions to change over time. Some have proposed studying microbial co-variations that are stable through time via conserved covariance, defined as the average of the binary covariance matrices across the observed time points after thresholding [8]. This is because microbial dynamics that are stable through time may provide information regarding the organization, structure and function of the microbiome. In particular, this paper shows that by considering the pairwise taxon-taxon associations in a microbial community, one can gain insights into conserved co-variation structure of the microbial community and its stability over time. However, since the relative abundance data are constrained to values between 0 and 1, and are often very sparse with many zeros, the standard measures of pairwise association such as Pearson’s correlation or rank-based correlations can lose power or lead to biased associations.

In this paper, we focus on the problem of estimating the conserved pairwise association between two bacterial taxon based on zero-enriched relative abundance data. Specifically, we define a general conserved dependence parameter between microbial pairs using generative copula models, with mixture margins, for the normalized relative abundance data. Copula models are advantageous because they separate the modeling of the dependence structure from that of the univariate margins. Such copula models have been shown to fit sparse microbiome proportion data well [9]. However, application of copula models to longitudinal data, outside of vine copulas, has been limited. We propose random-effects copula models where we assume that the dependence parameters across different time points follow a Gaussian distribution, with mean parameter that can be used to quantify the conserved microbial co-variations and to build conserved co-variation networks. This model assumes a random-effects model for the dependence parameter, which captures the conserved microbial co-variations, while allowing for a time-specific dependence parameter in the copula model.

To estimate the model parameters, we propose a Monte Carlo EM algorithm where Metropolis-Hastings sampling is used in the E-step. We also develop a Monte Carlo likelihood ratio test for hypothesis testing of the conserved dependence parameter. Our simulations show that the MCEM algorithm is efficient and provides an unbiased estimate the conserved dependence parameter in the random-effects copula models. We also show using simulations that the Monte Carlo likelihood ratio test has the correct Type I error. Finally, we present a detailed analysis of microbial co-variation networks in the infant gut microbiome and evaluate the effects of antibiotics on microbial co-variation network robustness, stability and centrality.

## Materials and methods

### Mixture Margin Random-effects Copula Models for Longitudinal Data

Consider a microbial sample taken from an individual at time *t*, that can be summarized by a vector of taxon counts: 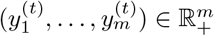. Due to sequencing constraints, these counts only contain information on the relative abundance of each taxon, thus making them difficult to compare across samples. For this reason, the counts are typically normalized by the total number of sequencing reads of the sample. That is, the normalized relative abundances of the *m*-microbes observed is defined as 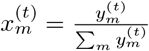 and denoted by the vector: 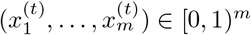.

Our focus is on measuring and testing co-variation among pairs of the microbes. Specifically, for any two of the microbes, *i* and *j*, let the joint cumulative distribution function of their relative abundances at measured at time *t* have the general copula form:

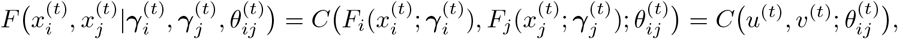

where 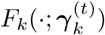 is the *k*-th univariate margin (*k* = *i, j*) with parameters 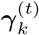 and *C*(·; *θ*^(*t*)^) is a family of copulas or multivariate uniform distributions with dependence parameter *θ*^(*t*)^, at time *t*. The univariate margins for microbial relative abundance data can be described by the zero-inflated beta distribution, which has proven to be a powerful and robust parametric distribution in the modeling of such data [10, 11]. The advantage of using the copula model is that the univariate marginal distribution allows us to include possible covariates 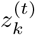 in modeling 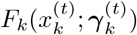, where the parameters 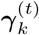 include the regression coefficients. For longitudinal microbiome studies, 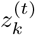 can include the relative abundance of the *k*th microbe observed at the the previous time point 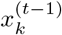, which leads the following auto-regressive marginal models:

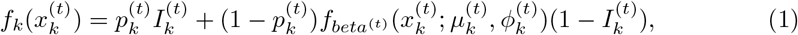

where we define 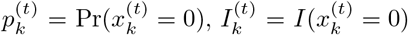, and

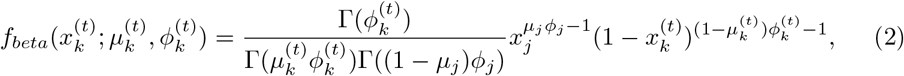

the density function of a beta random variable indexed by mean parameter 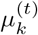 and dispersion parameter 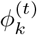. We can model 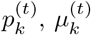, and 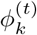 as a function of 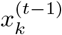 using logistic or log-linear regression models.

Henceforth, we omit the subscripts in the dependence parameter to ease notation, implying that we are referring to the modeling of a given (*i, j*) pair of microbes, unless otherwise noted. To define our proposed mixture margin random-effects copula models, we assume that the dependence parameter at time *t* follows a Gaussian distribution with the time-invariant dependence, denoted *θ*, between any two microbes, such that

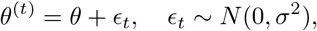

where *σ*^2^ is the variance parameter. This model allows for a time-specific dependence parameter *θ*_*t*_ between the two microbes, but assumes *θ* is the conserved dependence parameter that is used to measure the conserved co-variation between the two microbes. Larger variance *σ*^2^ corresponds to large variability of co-variations over time between the two microbes.

We are interested in estimating the parameter *θ* and in performing hypothesis testing of *H*_0_ : *θ* = *θ*_0_, where *θ*_0_ is some pre-specified, null value. In real microbiome applications, we test *H*_0_ : *θ*_*ij*_ = *θ*_0_ for each pair of microbes *i* and *j* and adjust for multiple comparisons by controlling for false discovery rate at a given level. We are most interested in the case where *θ*_0_ = 0 when Frank copula is used, which corresponds to independence between two microbes. We can then construct a co-variation network among all the microbes by creating links among those pairs with *θ*_*ij*_ ≠0.

### Joint density and the likelihood function

Given *θ*^(*t*)^, let the bivariate joint density function be given by 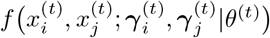, which can be be found via appropriate differentiation of the copula function *C*. When the margins are mixtures of absolutely continuous and discrete random variables, as is the case with the zero-inflated beta density, special care must be taken in the differentiation of the copula distribution function. A general framework for finding the density in such situations has been define, as well as the density for the special case of bivariate zero-inflated beta margins [9, 12].

The marginal likelihood at time *t* can be denoted as

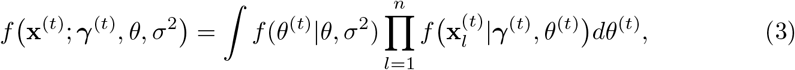

where 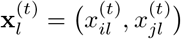 and 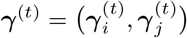. Additionally, assume the data from each of the *t* time points (*t* = 1, …, *T*) are conditionally independent and therefore the full data likelihood can be found by taking their product,

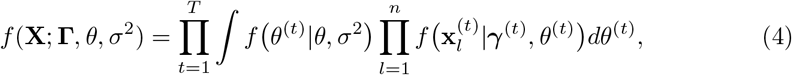

where **X** = (**x**^(1)^, …, **x**^(*T*)^) and **Γ** = (***γ***^(1)^, …, ***γ***^(*T*)^). It is important to note that the integration required in Eq. 4 is intractable. Thus, an approximate method for parameter estimation must be used.

### A two-stage estimation procedure

Under our proposed mixture margin random-effects model, the parameters of inferential interest are the true time-invariant dependence parameter *θ* and the variance, *σ*^2^, whereas the parameters of the time-dependent marginals, **Γ**, can be regarded as nuisance parameters. For ease of notation we define the full log-likelihood as

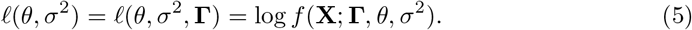

Consequently, we adopt a two-stage, or inference-for-margins, approach that is commonly used for inference involving copula models [13, 14]. The general two-stage estimation scheme can be described as follows:

1. Separately maximize each of the 2*T* univariate log-likelihoods,

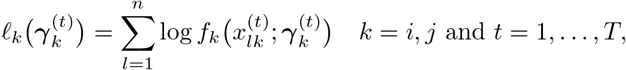

separately to get estimates of their parameters, 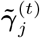 and 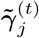, respectively.
2. Maximize the function 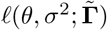 over *θ* and *σ*^2^ to get estimates 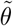 and 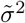.

Two-stage estimation reduces the computational complexity of the estimation procedure by replacing a single, joint optimization problem of many parameters with several smaller optimizations. However, step 2 of the algorithm cannot be solved directly due to the intractable integral in the likelihood function. Accordingly, we propose a Monte Carlo Expectation-Maximization (EM) algorithm to do so.

### A Monte Carlo Expectation-Maximization algorithm

The Monte Carlo EM algorithm [15] is particularly well-suited to solve the maximization of the marginal log-pseudo-likelihood, 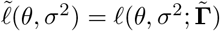. Under this framework, **X** are the observed data, *θ* and *σ*^2^ are the unknown, but fixed, parameters of interest, and **Θ**^(*T*)^ = (*θ*^(1)^, *θ*^(2)^, …, *θ*^(*T*)^) are unobserved latent variables or missing data. We define **(X, Θ**^(*T*)^) as the complete data and 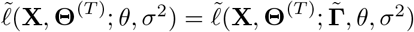 as its complete log-pseudo-likelihood function given by,

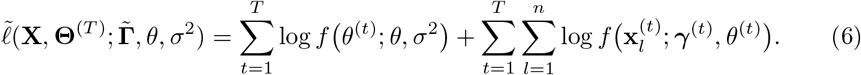

The assumption of the EM algorithm is that it is easier to maximize the complete data likelihood than the marginal likelihood. Given initial values 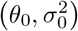, the algorithm produces a sequence of estimates, that converge to their incomplete data maximum likelihood estimates [16]. Let 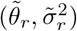 be the current estimates of *θ* and *σ*^2^ in the *r*^*th*^ EM iteration. The algorithm runs as follows for the (*r* + 1)^*th*^ iteration. First, in the expectation (E-) step, the missing data, **Θ**^(*T*)^, are replaced with their expectation:

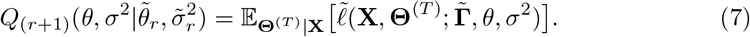

The expectation in (7) is taken with respect to the unobserved **Θ**^(*T*)^, conditional on the observed data **X** and the current estimates of the unknown parameters, 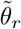 and 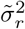. Based upon the definition of the complete data pseudo-likelihood in (6), 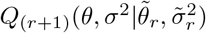 can be decomposed into two summations,

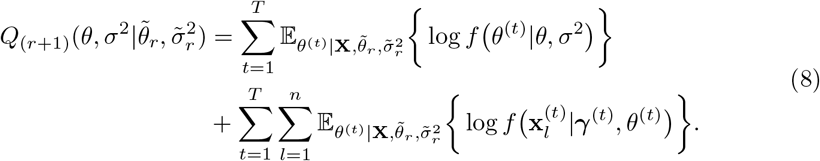

To ease notation, henceforth, we suppress the subscript and conditional terms of all expectations in the E- and M-steps. As such, expectations will be taken over the posterior distribution, 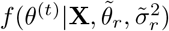, unless otherwise stated.

In the E-step, the expectation over *θ*^(*t*)^ cannot be directly computed, as the integration required to do so is intractable. We deal with the fact that we cannot directly compute these values by using a Markov Chain Monte Carlo (MCMC) sampling method, such as the Metropolis-Hastings algorithm. In the *r*^*th*^ iteration of the EM algorithm, we obtain a sample, 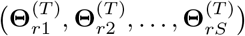, from 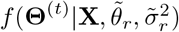, where *S* denotes the dependence on the MCMC sample size. Next, the M-step then maximizes *Q*_(*r*+1)_ to yield new estimates:

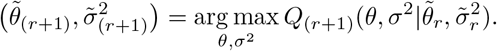

Partial differentiation of (8) with respect to *θ*_*r*_ and 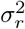 gives the following updated estimates,

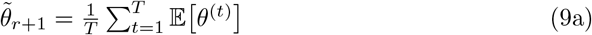

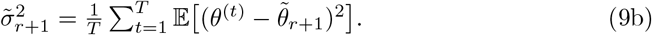

The above can be found by noting that the second summation in *Q*_(*r*+1)_ does not depend on either *θ* or *σ*^2^ and that each *θ*^(*t*)^ has density given by a Gaussian distribution with mean 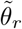 and variance 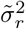 at the *r*^*th*^ iteration. The Monte Carlo estimates of the expectations are given by

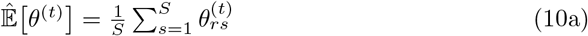

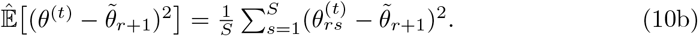

If the Markov chain converges to its target posterior distribution then, then 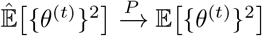 and 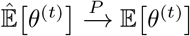.

Termination of the MCEM algorithm is not defined by usual convergence criteria used for the standard EM algorithm. This is due to stochastic nature of Monte Carlo portion of the algorithm. Instead, the average of the final *g* MCEM estimates are taken,

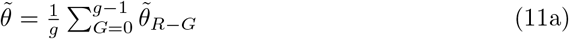

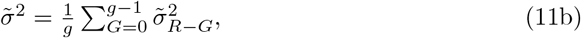

where *R* is the final iteration of the MCEM algorithm.

### A Monte Carlo likelihood ratio test

Suppose we are interested in determining if the two microbes have pre-specified dependence structure given by *θ*_0_. We can test this by comparing the likelihoods under our two-stage maximum likelihood estimate and the null parameter value. The likelihood ratio statistic is given by,

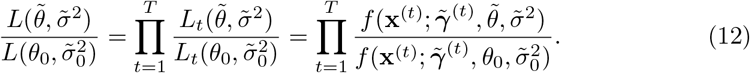

Typically, the log of (12) has an asymptotically scaled *χ*^2^ distribution [9]. As mentioned in the previous sections, direct calculation of these likelihoods are not feasible. Therefore, we use the following Monte Carlo approximation. Using the properties of conditional probabilities and Bayes rules, it has been shown [17] that Eq. 12 is equivalent to

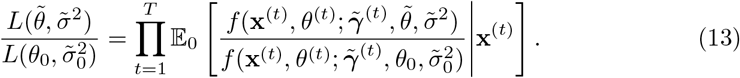

The expectation in Eq. 13 is taken with respect to the distribution of *θ*^(*t*)^ given **x**^(*t*)^ and the null parameters *θ*_0_ and 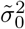. Finally, the Monte Carlo estimate of each of (13) can be formed using

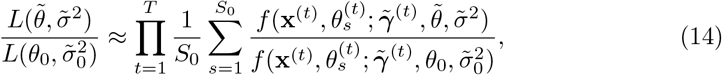

where 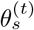 is a MCMC realization sampled from 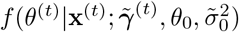. Such an estimation works the best when 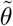 is close to *θ*_0_. We define the test using Eq. 14 as the Monte Carlo likelihood ratio test (mcLRT). Moreover, using the definition of conditional probability, the above ratio can be reduced to,

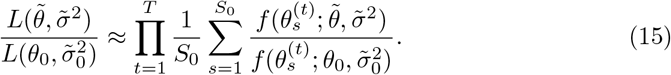

It is often the case that we are interested in testing for the independence of the two microbes, *θ*_0_ = 0 when the Frank copula is used, meaning the pair does not have a temporally conserved co-variation. In this setting, the asymptotic distribution reduces to a *χ*^2^ [9]. For the Frank copula, which is the parametric copula function we focus on, *θ*_0_ = 0.

## Results

### Simulation Studies

Simulation studies were used to assess the estimation accuracy of the two-stage Monte Carlo EM procedure. The data was simulated according to the following generative model, using the Rosenblatt transformation of the Frank copula function. We focus on the Frank copula function as it can model both positive and negative dependence, and the magnitude of dependence is symmetric. First, define *U* = *F*_*i*_(*x*_*i*_), *V* = *F*_*j*_(*x*_*j*_) and

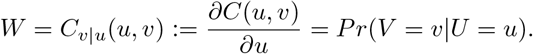

The generative algorithm proceeds as follows:

1. Set *θ* and *σ*^2^
2. For every *t* = 1, …, *T* and *l* = 1, …, *n*
  a. Draw *θ*^(*t*)^ ∼ *N* (*θ, σ*^2^)
  b. Draw 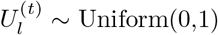 and 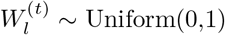
  c. Solve for 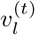 using:

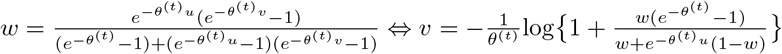
  d. Solve for 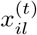 using the definition of *U* :

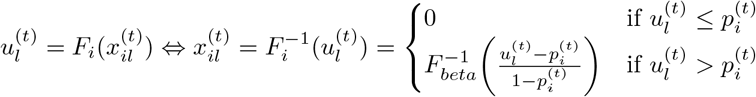

Likewise, the procedure for 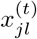 and 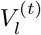 is the same.

The above process is repeated if (i) if there are less than three non-zero relative abundances for either microbial taxon, as this is the minimum number non-zero observations needed to estimate the three marginal parameters, (ii) if the two taxa are mutually exclusive, i.e. if one taxon has a non-zero relative abundance the other must be absent, or if only one pair of observations has non-zero relative abundance for both taxa. This avoids the situation where the dependence parameter hits the lower boundary of estimation and/or causes unstable variance estimates.

Simulations were performed under varying parameters values for *θ* and *σ*^2^ to evaluate the robustness of the estimation procedure. Three simulation scenarios assumed intercept-only, or no covariate effect, models for the margins. For these models the zero-inflation probabilities are set to (*p*_*i*_, *p*_*j*_) = (0.4, 0.5). Additionally, the mean and dispersion parameters of the beta portion are set to (*μ*_*i*_, *μ*_*j*_) = (2*/*7, 3*/*9) and (*φ*_*i*_, *φ*_*j*_) = (7, 9), respectively. Finally, the true dependence parameter was selected as either *θ* = 0.5 or 4 and standard deviation *σ* = 0.5 or 1.3.

Since the procedure estimates the marginal parameters independently, we can incorporate dependence between successive relative abundances by using an autoregressive model. Under this framework, an AR(*q*) model includes the relative abundance from the previous *q* time points as covariates in regression model for *p, μ* and/or *φ*. We simulated data from an AR(1) structure for the model for the zero-inflation parameter and model for the mean. Furthermore, we set the true parameter values to be as follows: (***ρ***_*i*_, ***ρ***_*j*_) = ({1, −4}^T^, {1, −4}^T^), (***δ***_*i*_, ***δ***_*j*_) = ({−1, 1.5 }^T^, {−1, 1.5} ^T^), (*κ*_*i*_, *κ*_*j*_) = (1, 1), *θ* = −3, and *σ* = 1.

Simulations were repeated fifty times for each of the four model settings, the sample size for each was set to *n* = 100, and the number of time points to *T* = 25. Moreover, for each simulated data set, the EM algorithm was run for 20 iterations and the Markov chains were run for a length of 4000. Figure 1 shows that, in general across the fifty simulation runs, 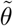 is an unbiased estimator. Similarly, our 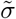 estimator is also almost unbiased. Furthermore, the trace plots of the EM estimates at each iteration (not shown) show random variability of the estimates around the true value. These plots also show that the MCEM algorithm quickly stabilizes for the 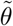 estimates, while that of the 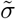 estimates is slower.

**Fig 1.**
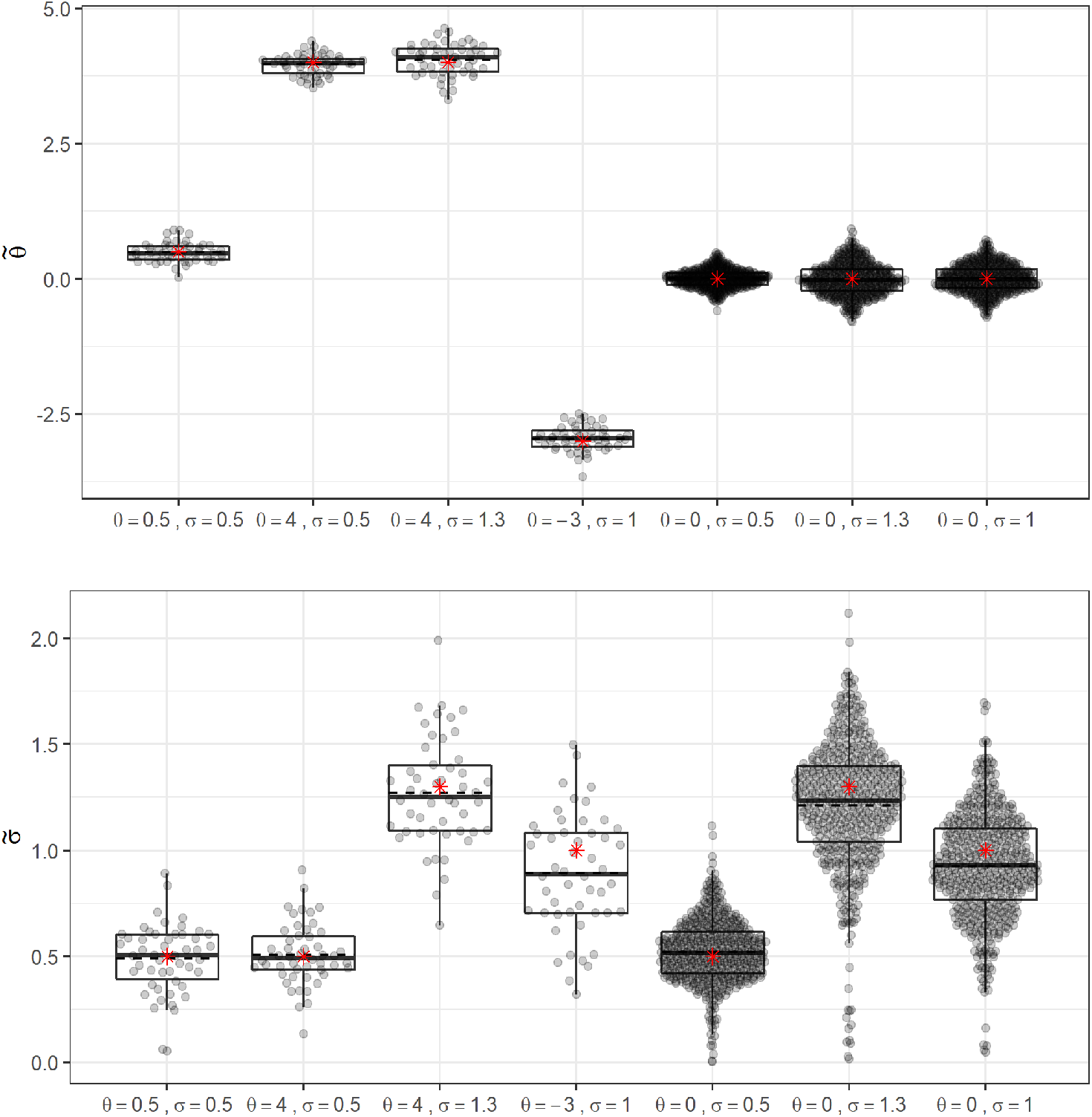
Box plot of the 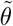 and 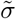 estimates under seven different simulation settings (null and alternative). The black dashed line represents the mean across all runs. The red star indicates the true value set in simulation.

We further use simulations to assess the Type I error rate of the Monte Carlo likelihood ratio test for independence. To do so, data was simulated under the independence model (*θ* = 0). The remaining model parameters were specified as above, as well as the sample size and number of time points. The EM algorithm was run for 20 iterations and the Markov chains had a length between 2500-4000. The estimation results from 250 replicates were similar to the those above, including almost unbiased MCEM estimate 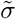 for large values (Figure 1). Finally, the Type I error rate of the Monte Carlo LRT is 0.058 and 0.052 for the AR(0) models with *σ* = 0.05 and *σ* = 1.3, respectively. For the AR(1) model with *σ* = 1 the Type I error is 0.066. Thus, we concluded the Type I error rate is controlled at the nominal 0.05 level.

### Analysis of Temporally Conserved Microbial Networks During Childhood Development

#### Microbial co-variation network estimation in children with and without antibiotic exposure

We applied this method to the DIABIMMUNE antibiotics cohort from the Broad Institute [18]. The cohort consists of 39 children from Finland. The gut microbiome of each child is densely sampled with an average of 28 samples per child collected monthly over the first 36 months after birth, thus making the data set particularly well suited for our method. These data were originally used to perform a natural history study to explore the dynamics of the gut microbiome of children before stabilization to a mature state [18]. The data is publicly available under NCBI BioProject ID PRJNA290381. In this analysis, we used the bacterial relative abundance data from the 16S sequencing.

Twenty children received between nine and fifteen antibiotic treatments, thus allowing quantification of the effects of antibiotic use in comparison to those who had never received antibiotics. As a result we split the data set into two groups (antibiotics vs. no antibiotics) and performed estimation and testing of conserved co-variation measures between microbe pairs. The study collected a total of 1002 samples, 483 of which are from children who received antibiotics and 519 from those who did not. We removed any sequencing reads that were unassigned or ambiguously assigned at the genus level. Furthermore, we only used genera with at least a 10% prevalence across all samples in each group. This resulted in a total of 51 unique genera, from which 1275 pairs can be formed. For each pair, time points with less than 25% prevalence for either taxon were removed and any pair with less than ten time points remaining were removed. This left a total of 885 and 878 pairs for antibiotic and no antibiotic groups, respectively. For all pairs, we performed MCEM estimation and testing for independence between the two microbes. We adjusted for multiple comparisons by controlling the false discovery rate at 5% using the Benjamini-Yekutieli procedure [19].

A total of 61 and 110 pairs had a significant dependence parameter after FDR control in the antibiotics and no antibiotics groups, respectively. These significant pairs were then use to construct the conserved co-variation networks of the microbial community in children with or without antibiotic exposures. To illustrate the conserved nature of these dependency structures, Figure 2 plots the posterior mean of the Monte Carlo sampled time-dependent dependence parameters (i.e. *θ*^(*t*)^) for a small subset of pairs from each network. Pairs with a small dependence parameter have little variability through time, whereas such variability grows for larger dependence parameters, particularly negative ones, but are still centered around the conserved estimate.

**Fig 2.**
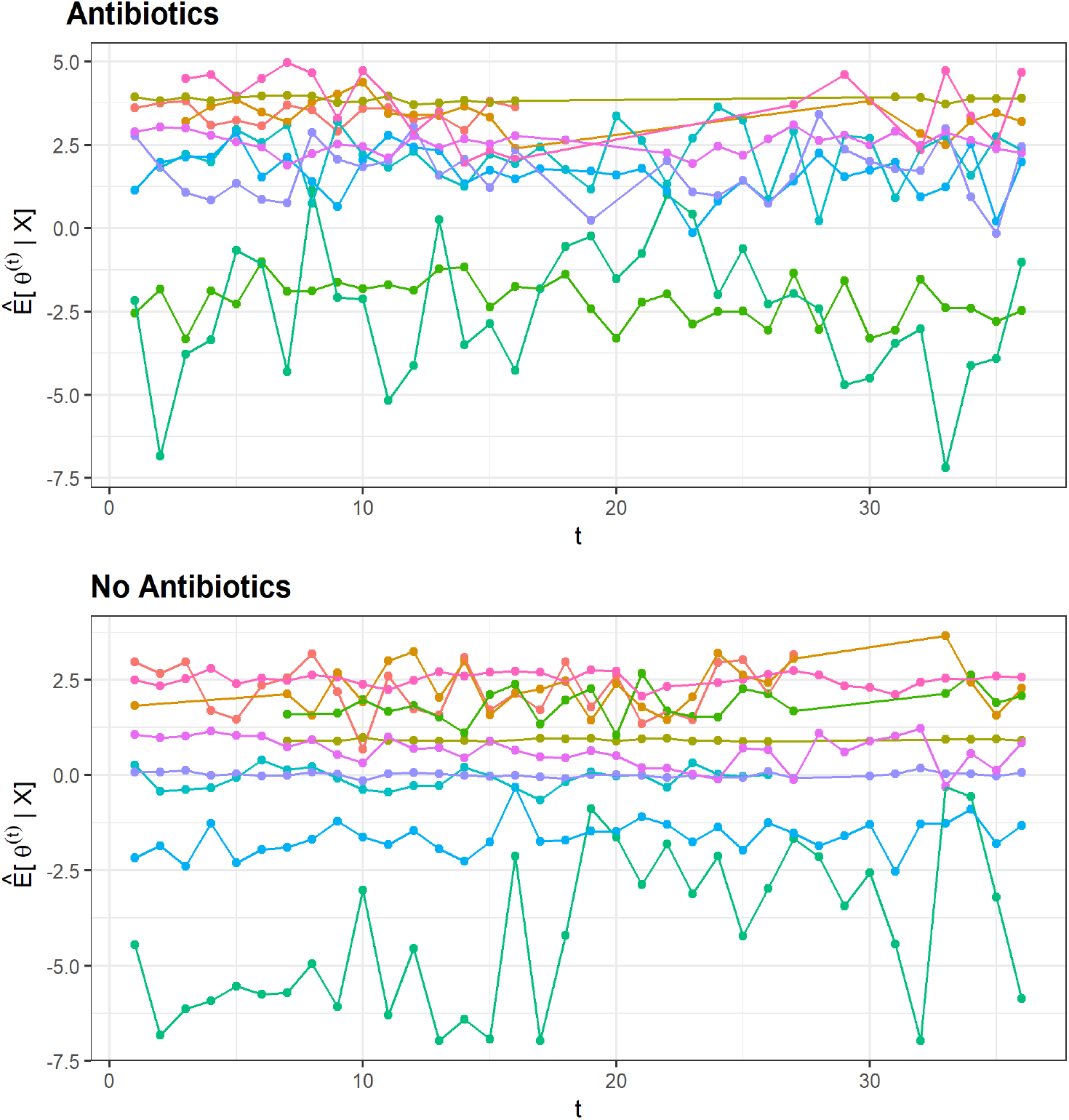
Plot of the posterior mean of *θ*^(*t*)^ for a random subset of ten significant pairs in each network, showing relatively constant dependence parameters over time.

#### Comparison of co-variation network properties

From these results we built an adjacency matrix, **A**, and performed co-variation network analysis. Two microbes are said to be connected (*a*_*ij*_ = 1) if their dependence parameter was significant and are unconnected (*a*_*ij*_ = 0) otherwise. Each of the 51 genera are represented by the nodes of the network diagram and edges correspond to the entries of the adjacency matrix (Figure 3). The graphs show that the overall network structures differ between the two groups, with only 29 common edges. The antibiotics group is less densely connected than the no antibiotics network, as seen by its higher mean distance (3.32 vs. 3.02), lower edge density (0.048 vs. 0.086), and lower cluster coefficient (0.260 vs. 0.377). Both networks have five clusters, and consist of several unclustered taxa, as identified from a fast-greedy algorithm that aims to maximum network modularity [20]. Additionally, the antibiotics network is more modular (0.539 vs. 0.302) with many connections within clusters and few between. The adjusted Rand index can be used to quantify the similarity between the clusters detected in the two networks. We observe the adjusted Rand index between the antibiotics and no antibiotics groups to be 0.132, as compared to 0.023 ± 0.018 obtained from permutation testing, indicating the higher overlap between the two is more than expected by chance. Finally, we detect there to be a significance difference (*p <* 0.001) between four-node motif occurrence in the two networks, indicating there is also a difference in their local organizations.

**Fig 3.**
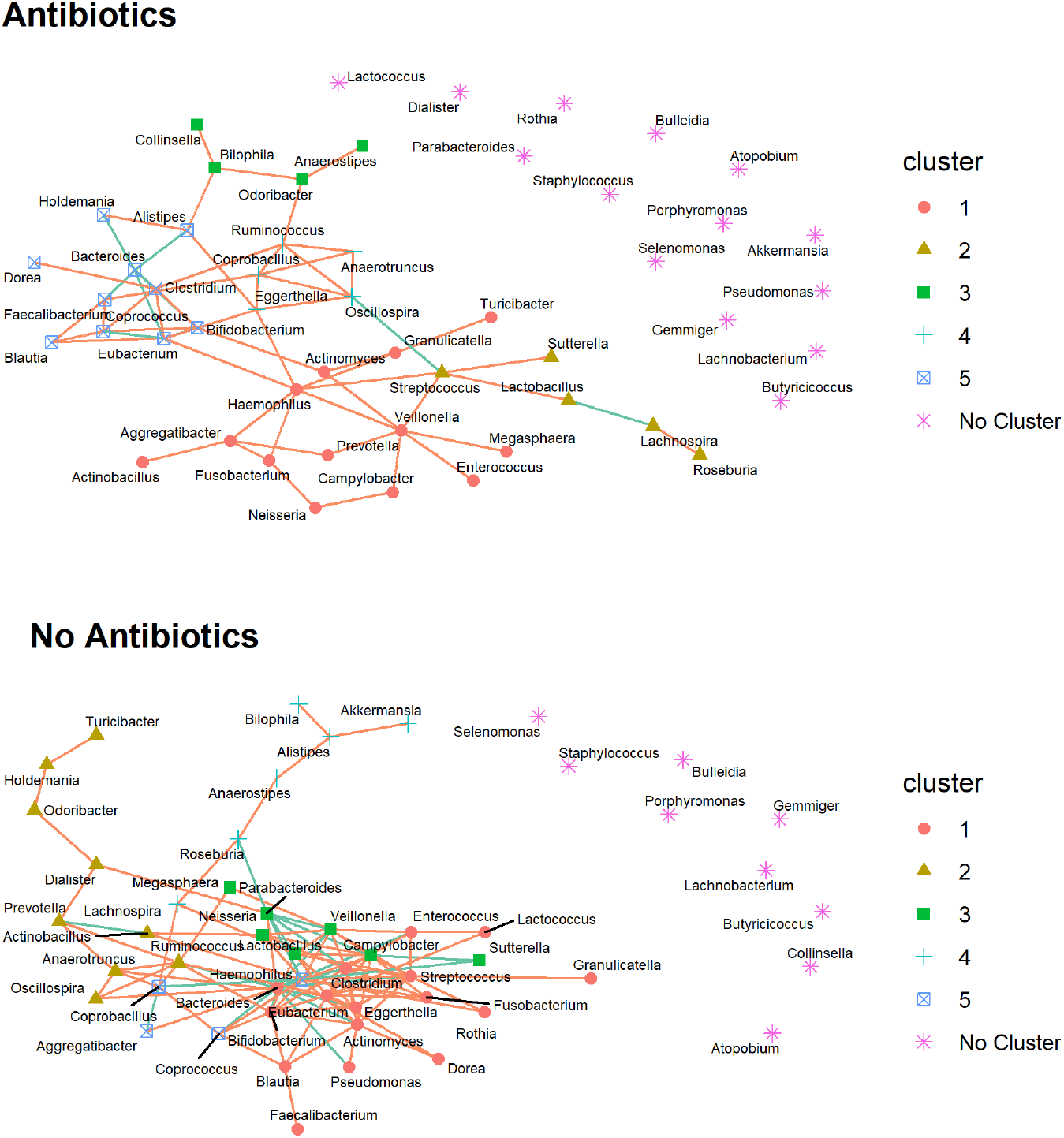
Conserved microbial co-variation network diagrams for the antibiotics and no antibiotics groups, respectively. Each node represents a bacterial genus and its shape and color correspond to graph cluster from an algorithm that maximized the modularity. Edge color corresponds to positive/negative (orange/green) dependence.

We further compared the two networks using node-specific centrality parameters. The antibiotics network has an average degree of 2.39 (sd=2.22), average closeness of 0.049 (sd=0.018), and average betweenness of 0.025 (sd=0.041). In contrast, the no antibiotics network has an average degree of 4.31 (sd=4.19), closeness of 0.074 (sd=0.026), and betweenness of 0.028 (sd=0.042). The lower average degree, closeness and betweenness of the antibiotics network compared to that of the no antibiotics provides further evidence of a less connected network. Figure 4 shows that the majority of microbes have a higher degree and closeness in the no antibiotics network than that of the antibiotics network.

**Fig 4.**
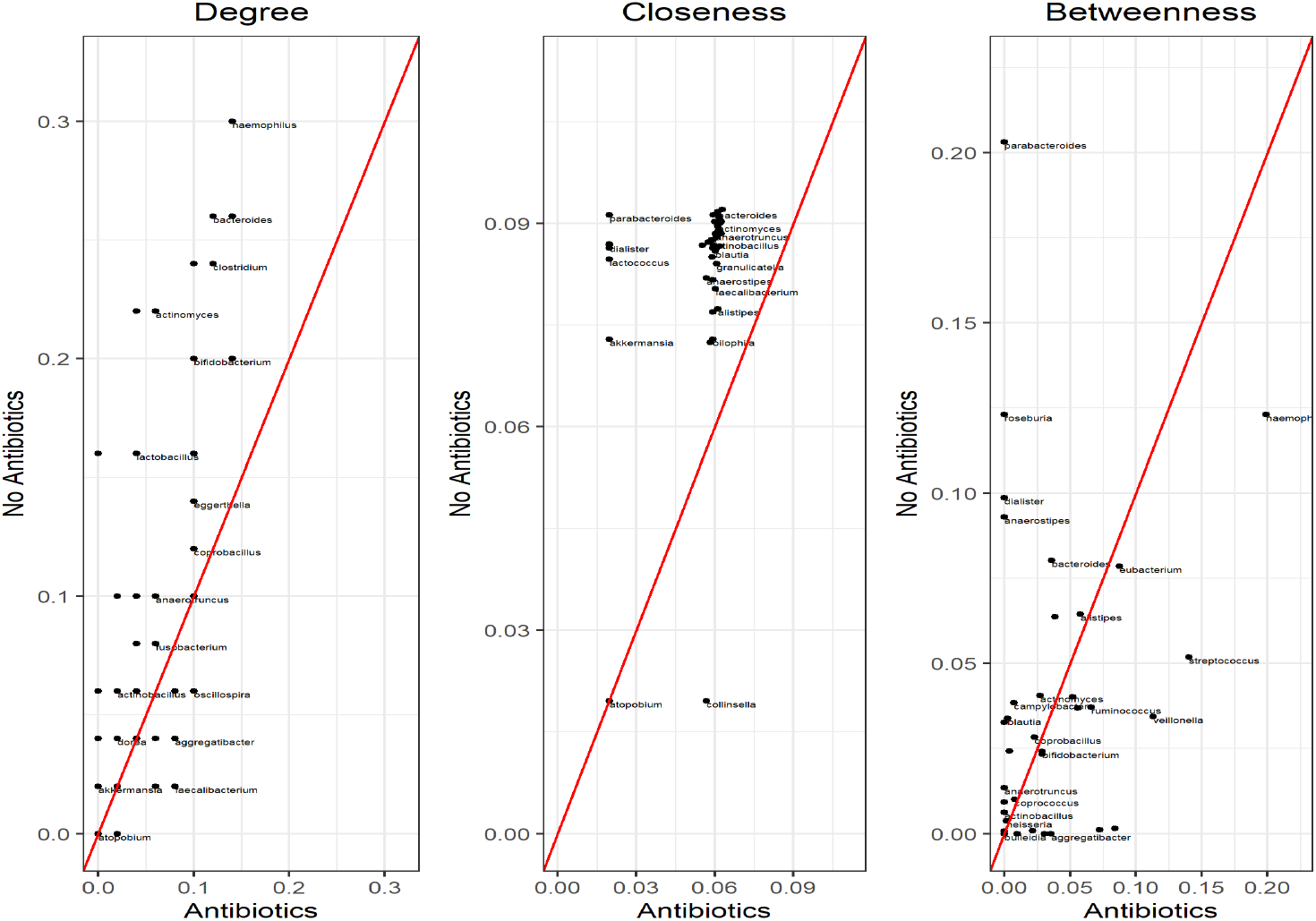
Node-wise centrality statistics for each microbe in both the antibiotics and no antibiotics networks. Red line corresponds to *y* = *x*.

#### Network Robustness and Stability

We also assessed the robustness and stability of the networks using targeted and random attacks (Figure 5). The ‘attacks’ involve sequentially removing the nodes from the network. Targeted attacks remove nodes in decreasing order of degree (i.e. nodes with higher degree are removed first) and random attacks remove nodes with equal probability. For each removed node, the diameter, defined as the longest path, and global efficiency, the average of the shortest path between all pairs, are calculated. Targeted attacks show that it takes until about 10% of the nodes are removed for the diameter to change in the no antibiotics network, for subsequent nodes the two networks follow similar trajectories with that of the no antibiotics network shift forward by five to ten nodes. The efficiency of the two networks under targeted attacks show a similar decreasing pattern starting with the first node removal (results not shown), which is to be expected as we are removing the most connected nodes first. Despite showing a similar pattern the efficiency of the no antibiotics network remains uniformly more powerful than that of the antibiotics network. Under random attacks, the efficiency of the no antibiotics network remains larger than that of the antibiotics networks until about 40% of the nodes are removed, at which point it drops to the same level as the antibiotics network. Together these results indicate that the no antibiotics network is less dependent on individual nodes, thus making it more stable than the antibiotics network, and is more robust to both targeted and random attacks.

**Fig 5.**
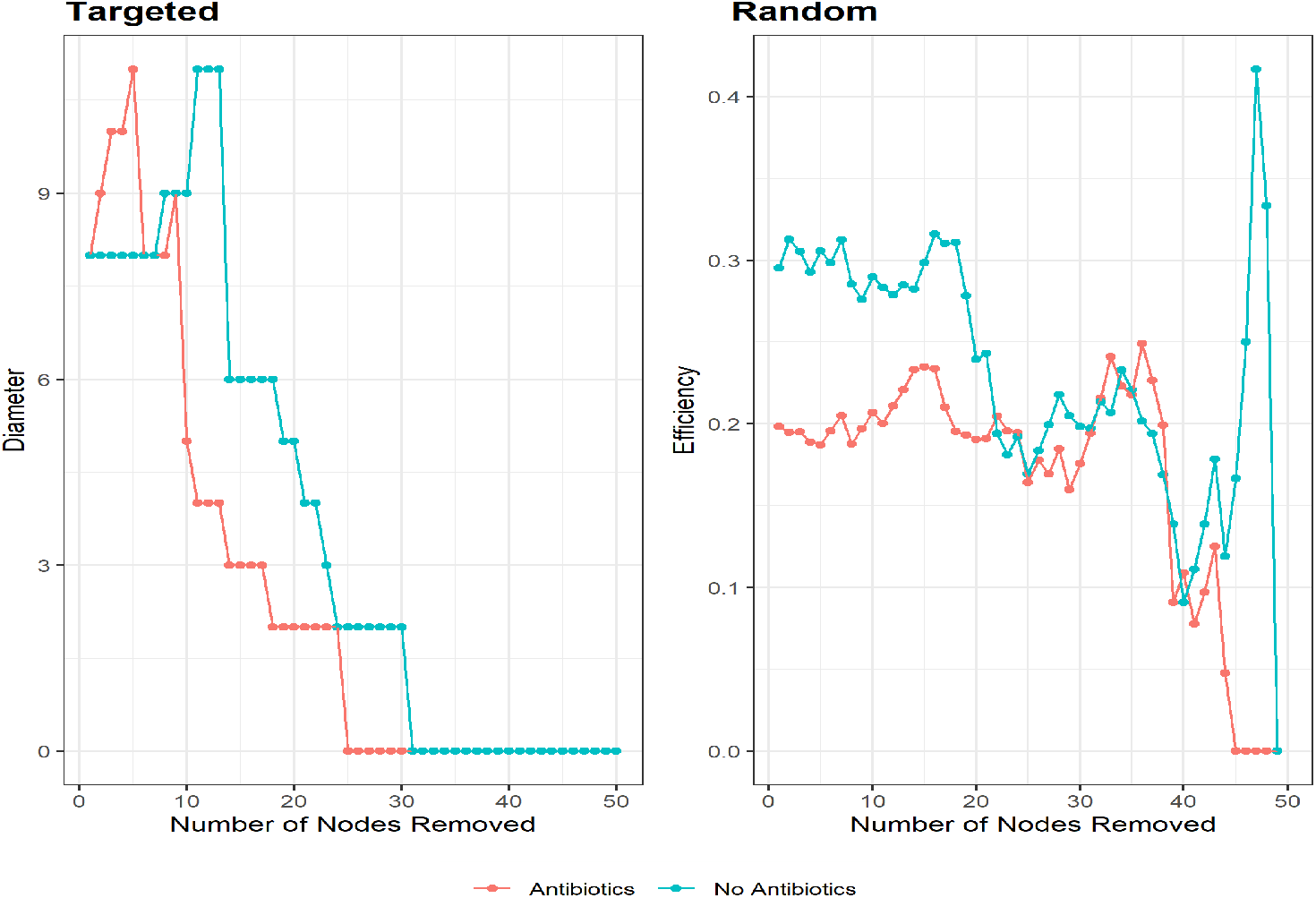
Network fragility as measured by diameter and efficiency in targeted and random attacks, respectively.

## Discussion

To fully understand the functional properties and organization of the microbiome, it is necessary to begin studying its emergent properties. Such properties cannot be identified from single species, “parts-of-a-whole” approaches, as they only appear from interactions between the parts of the system. We define the bivariate dependency structure between all pairs of taxa as the emergent properties of interest in this paper. In particular, we focus on those dependence structures that are temporally conserved to understand the organizational stability of microbiomes.

To identify such dependence patterns, we have developed a mixture margin random-effects copula model by assuming the time-specific dependence parameter of bivariate copulas follow a Gaussian distribution. At each time point the observed data are modeled with a bivariate copula distribution, with a time-specific dependence parameter. Each of the time-sensitive parameters as assumed to be a random observation from a Gaussian distribution whose mean captures the conserved dependence structure. We derived an efficient Monte Carlo EM algorithm for estimation of the time-invariant dependence parameter and variance parameter, as well as a corresponding Monte Carlo likelihood ratio test. Our method is applied to real data to understand the impact of antibiotic use on the organizational structure of the infant microbiome. We find that antibiotic use leads to both global and local changes in the resulting microbial networks of conserved co-variations. Furthermore, antibiotic use is associated with a network that is less robust when subjected to both targeted and random attacks.

We have focused on the estimation of pairwise co-variations as they are useful when the sample size is small, as is the case in the DIABIMMUNE data. The proposed model can be used to assess the conditional pairwise associations by including all other taxa in the marginal models. Our model is developed to assess the conserved co-variations between two taxa across time. As a future research, it is interesting to extend the methods to allow for change-points in co-variations over time.

## Conclusion

The proposed mixture margin random-effects copula models are shown to be able to capture the conserved co-variation relationship between two bacterial taxa based on longitudinal microbiome data. Such co-variation relationships among the bacteria lead to construction of conserved co-variation networks. Comparing such networks between two biological conditions can provide insights into how environmental factors, such as antibiotics use, affect the network structures. Using the DIABIMMUNE longitudinal data, we have shown that the majority of microbes have a higher degree and closeness in the no antibiotics network than that of the antibiotics network. In addition, the no antibiotics network remains uniformly more powerful than that of the antibiotics network under targeted attacks. Under random attacks, the efficiency of the no antibiotics network remains larger than that of the antibiotics networks until about 40% of the nodes are removed, at which point it drops to the same level as the antibiotics network. Together these results indicate that the no antibiotics network is less dependent on individual nodes, thus making it more stable than the antibiotics network, and is more robust to both targeted and random attacks.

## Acknowledgments

This research is supported by NIH grants GM129781 and GM123056. *Conflict of Interest*: None declared.

## Notes

### Competing Interest Statement

The authors have declared no competing interest.

